# Cellector: A tool to detect foreign genotype cells in scRNAseq data with applications in leukemia and microchimerism

**DOI:** 10.64898/2026.03.26.714571

**Authors:** Haynes Heaton, Reza Behboudi, Collin Ward, Minindu Weerakoon, Sami Kanaan, Skylar Reichle, Nathan Hunter, Scott Furlan

**Affiliations:** Auburn University, Auburn, AL 36849; Fred Hutchison Cancer Center; Seattle Childrens Hospital, Department of Pediatrics

## Abstract

The existence of rare, genetically distinct cells can occur in various samples such as transplant patients, naturally occurring microchimerism between maternal and fetal tissues, and cancer samples with sufficient mutational burden. Computational methods for detecting these foreign cells are vital to studying these biological conditions. An application that is of particular interest is that of leukemia patients post hematopoietic cell transplant (HCT). In many leukemias, a primary therapy is HCT, after which, the primary genotype of the bone marrow and blood cells should be of donor origin. If cells exist that are of the patient’s genotype and the cell type lineage of the particular leukemia, this is known as measurable residual disease (MRD). If the MRD is high enough, this may represent a relapse of the patient’s leukemia. Furthermore, accurately estimating the MRD is important for driving clinical decision making for these patients. Here we present Cellector, a computational method for identifying rare foreign genotype cells in single cell RNAseq (scRNAseq) datasets. We show cellector accurately detects microchimeric cells down to an exceedingly low percentage of these cells present (0.05% or lower).

---

The ability to detect exceptionally rare, genetically distinct, cells is important for the study of microchimerism, transplant biology, as well as early detection of leukemia relapse. Using high throughput single cell RNAseq (scRNA-seq) such as drop-seq^1^, 10x Genomics^2^, Seq-Well^3^, InDrops^4^ among others, one can use the expressed genetic variants in the RNAseq reads to detect microchimeric cells. Unlike multiplexed single cell experiments, one cannot biochemically tag these cells for demultiplexing^5,6^. Tools made for demultiplexing cells by genotype such as souporcell^7,8^, vireo^9^, and scSplit^10^ rely on clustering based systems that don’t perform well with highly skewed cluster sizes, which microchimerism has by definition. Other tools such as Demuxlet^11^ require knowledge of the genotypes up front which may be costly, not possible, or unavailable. Here we present Cellector, a tool to accurately detect microchimeric cells while being specific down to exceedingly low numbers of cells. Cellector uses a sparse beta-binomial anomaly detection method to identify cells with different genotypes than the majority of the sample. We show that cellector can accurately identify foreign cells down to 0.05% or lower even when the cells come from related individuals which represent the most common donors for HCT. Cellector is freely available under an MIT open-source license at https://github.com/wheaton5/cellector.

## Results

### Genetic anomaly detection and data processing

In order to detect genetically distinct cells, we must first detect which alleles of each genetic variant are expressed in the RNAseq reads of each cell. To achieve this, we find expressed regions overlapping common human variants occurring at least 1% in the population according to population scale genome projects ^12,13^. We then count alleles for each cell barcode using vartrix (Fig. 1a)^14^. We then create a beta-binomial distribution for each genetic variant with parameters alpha and beta being the total number of alleles expressed for the alternative and reference alleles respectively (with 1 added to these parameters for the bayesian conjugate prior/uniform beta distribution) (Fig. 1b). We treat these distributions as the distribution of alleles for the majority genotype individual in the sample. We can then compute a log likelihood of the alleles expressed in each cell given these distributions (treating each loci independently) and normalize this value for the fact that some cells will express many more reads covering genetic variants (Fig. 1b,c). We can then use anomaly detection on this normalized log likelihood value to find foreign genotype cells (Fig. 1d). We then iteratively remove those anomalous cells’ alleles from the beta-binomial distribution parameters for those variants, and repeat anomaly detection. We continue this process until we converge on one set of cells being detected as genetically distinct (Fig. 1e). Finally, we create two beta-binomial distributions for each locus, one for the majority cells, and one for the minority cells. Using these two distributions, we can calculate a posterior probability for each cell to those distributions for a final call of minority cells (Fig. 1f).

**Fig. 1.**
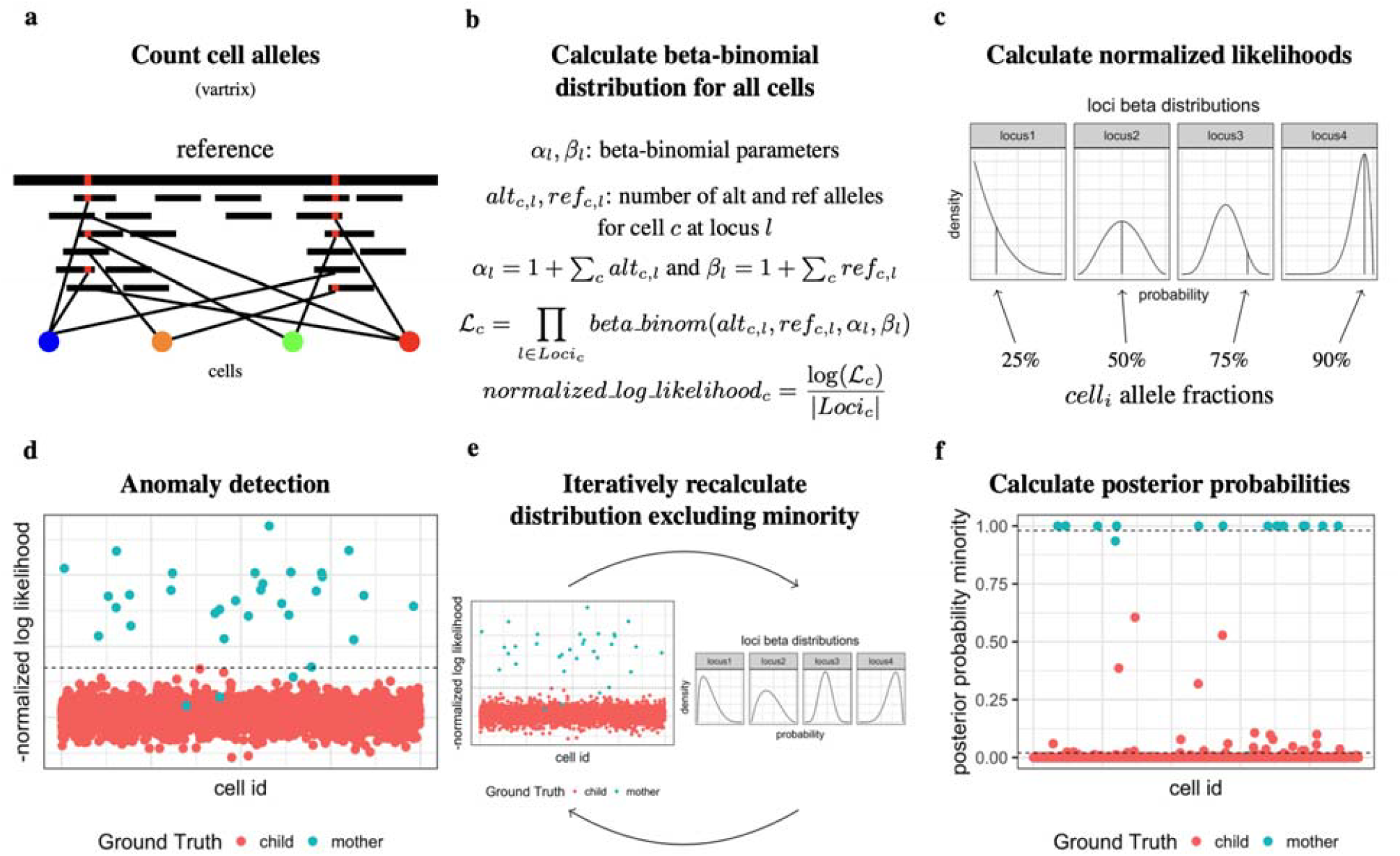
Cellector overview. **a**, We first count alleles for each cell for expressed regions for common variant loci. **b**, We then create beta-binomial models for each of these loci for all cells. **c**, We calculate the normalized log likelihoods of each cell given their expressed alleles. **d**, Run anomaly detection to identify genetically distinct cell. **e**, Iteratively remove allele counts from majority genotype distribution and recall anomalous cells until convergence. **f**, Create distributions for majority cells and minority cells and calculate posterior probability assignments.

Cellector makes the important assumptions that a majority of the cells in a sample are from one genotype and there may or may not be a minority (<20%) of cells that are from foreign genotype(s). These assumptions give us an advantage over other methods such as clustering by genotype where the number of individuals could be any number and the clusters could be of any size. Using anomaly detection is vastly more sensitive and specific than clustering when the number of minority genotype cells is very low. Because of these assumptions, if the minority is not less than 20%, Cellector starts losing sensitivity because 20% of a dataset isn’t that anomalous. But in these cases, clustering by genotype works well. To overcome this issue, we run both Cellector and our previous tool, souporcell, independently and choose the output according to how well each tool separated the allele distributions. To measure this, we create beta-binomial distributions for majority and minority according to assignments of each program and calculate the mean difference in log likelihoods for the two different distributions across all cells. Whichever program creates a larger log-likelihood difference is chosen as better separating the minority and majority distributions.

### Validation and benchmarking

We validated Cellector against both real mixtures with cell hashing and also in-silico mixtures for full ground truth and the ability to test against a range of conditions. We focused on related individuals to validate for the application of leukemia relapse post HCT. We made cell hashing mixtures for two families with three different mixture levels for each. For the first, we have a haploidentical mother and child with 250, 50, and 10 maternal cells spiked into 10,000 cells each from the child (Fig. 2b). These samples recovered far fewer than the targeted 10,000 cells with ∼2750 cells recovered in each run.

**Fig. 2:**
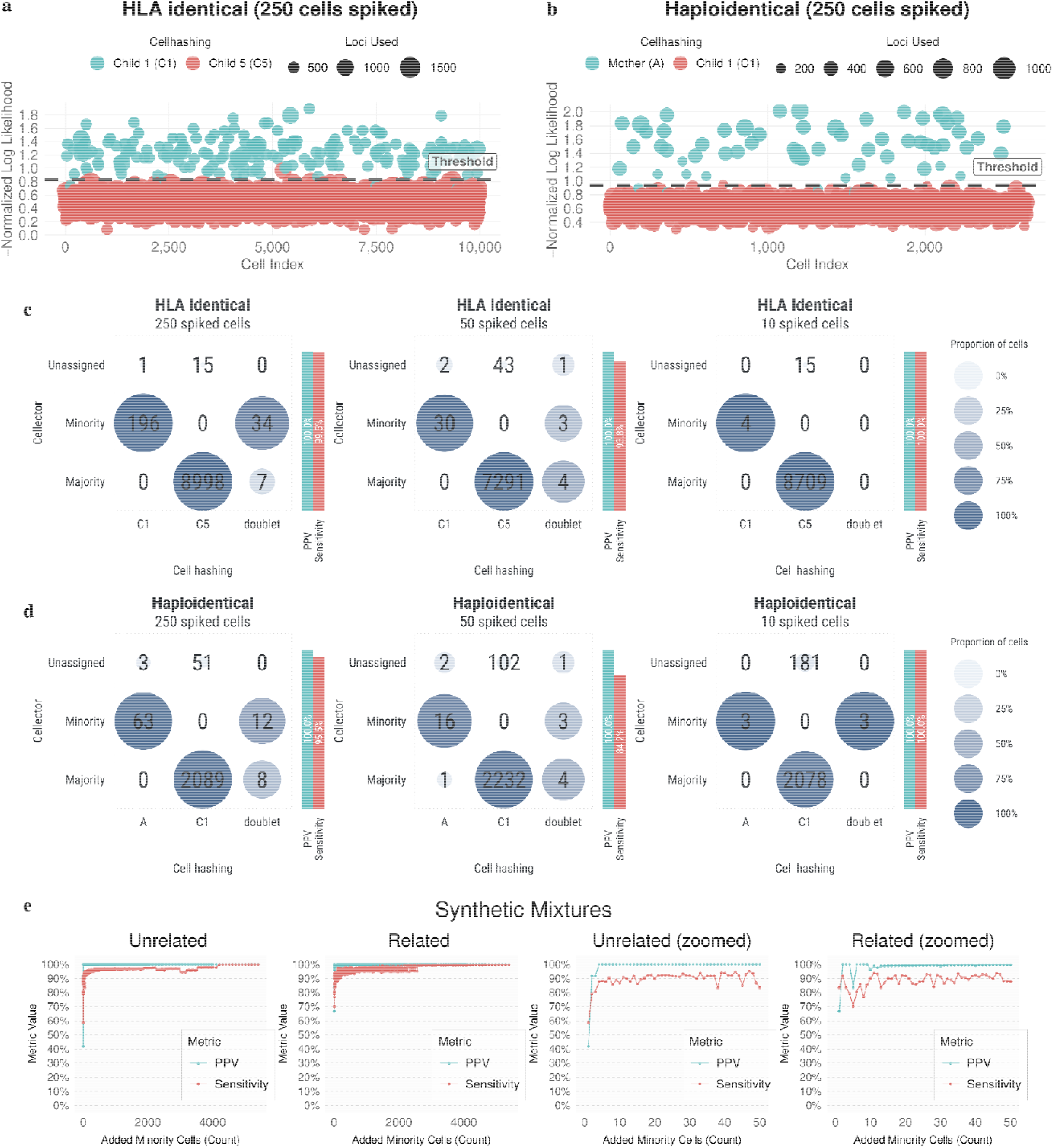
Cell hashing and in-silico mixture validation. **a**, Anomaly plot for cellector on a mixture of HLA-identical sibling cells with 250 minority cells spiked into 10k cells with cell hashing. **b**, Anomaly plot for cellector on a mixture of haploidentical parent/child cells with 250 minority cells spiked into 10k cells with cell hashing. **c**, Comparison of cellector to cell hashing for mixtures of HLA-identical siblings with 250, 50, and 10 cells spiked into 10k cells. **d**, Comparison of cellector to cell hashing for mixtures of haploidentical parent/child samples with 250, 50, and 10 cells spiked into 10k cells. **e**, Sensitivity and PPV across a range of minority-cell fractions for in-silico mixtures of unrelated and related individuals.

You can see clear separation of minority cells versus majority cells in our anomaly plot (Fig. 2b) with the few minority below the threshold often be saved in the posterior probability step. We compared Cellector’s assignments to the assignments from cell hashing (Fig. 2d). We exclude doublets from our sensitivity and PPV numbers because cell hashing drastically overcalls doublets and the expected number of doublets is very low. We show perfect PPV while maintaining sensitivities between 93% and 100% and we leave ∼3.5% of cells unassigned. Cellector misidentified one minority cell according to cell hashing as a majority genotype cell (Fig. 2d middle panel). We even detected all 3 cells identified as maternal by cell hashing for the spike-in of 10 in 10k (Fig. 2d, right panel).

For the next cell hashing validation, we collected cells from two HLA identical brothers. These experiments yielded more cells (∼8k to 10k) which is useful to test Cellector under a higher doublet barcode setting. We again see good separation on the anomaly plot (Fig. 2a), perfect PPV with sensitivities between 84% and 100%. We again detect all 4 cells from the spike in mixture of 10 in 10k cells (Fig. 2c right panel).

In addition to the cell hashing validation, we wanted to test Cellector’s performance across a whole range of spike in levels while also having real ground truth as opposed to the sometimes faulty cell hashing (especially faulty in doublet calling). So we created in-silico mixtures of separate scRNAseq datasets such that we could control how many cells were mixed as well as have perfect ground truth. We did this with both related individuals and unrelated individuals. We obtained cells from a family with both grandparents, a mother, and two children. We synthetically spiked in cells from one sample into another sample. We ran these experiments across a whole range of spike in levels and averaged across multiple random runs across all direct-relation pairs. We tracked the sensitivity and PPV for these runs (Fig. 2e,g) as we swept the number of cells being spiked in. Cellector almost always maintained a PPV of 100% with a few mistakes across thousands of runs. The sensitivity was generally above 90% except with very few cells spiked in it dropping to 70%. We see a similar picture with unrelated individuals which were taken from the human induced pluripotent stem cell project (HipSci) (Fig. 2f,h) except that PPV was perfect with sensitivity again being above 90% except when very few cells are present.

### Comparison to existing tools

While limited tools exist for this application, some can work in certain conditions. So here we compare the performance of Cellector versus other scRNAseq demultiplexing software including our previous tool, souporcell, vireo, and demuxlet across # of cells spiked in in-silico mixtures of unrelated (Fig 3 a,b) and related (Fig 3 c,d). We show that cellector maintains high (>90%, often perfect) PPV down to even a single cell whereas other demultiplexing tools have low PPV until ∼1.5-3.5% minority cells. Most tools have fairly high sensitivity in these low ranges in exchange for low PPV. For clinical and research applications, it is vital to have high confidence in the assignment of these foreign cells. We did not have genotype information for the related individuals and thus could not compare demuxlet to other tools in those samples. We also stress tested Cellector in terms of the amount of data the samples had by running a series of downsampling experiments with 10 and 100 foreign cells added (Supp Fig 2) and show that Cellector maintains near perfect PPV until median UMI per cell drops below 904 while maintaining reasonably high (>85%) sensitivity. In these experiments, we compared to souporcell which does not find the optimal clustering with just 10 cells, but with 100 cells tends to trade off lower PPV for higher sensitivity and its performance drops off more the less data it has.

### Additional applications and sample types

Here we deviate from our main focus on the Leukemia application an show results of Cellector being applied to maternal/fetal cell atlas samples^15^, a neuronal sample with known microchimerism^16^, and kidney transplant samples. In samples from the maternal/fetal cell atlas^15^ we show that cellector is able to identify cells in placental samples that are of maternal origin. In these samples, it is expected that the cell types are different between maternal cells and fetal cells which are corroborated by the t-SNE (Fig 4 a,b). The maternal origin cell types correspond to maternal macrophages and maternal decidual stromal cells known to be an experimental contamination in these samples. We then show cellector’s ability to detect microchimeric cells in a neuronal sample (Fig 4 c)^16^. In this sample, cells are not expected to be of different cell types due to the hypothesis that these cells migrated to the brain while still pluripotent and differentiated according to the normal tissue microenvironment. But in the anomaly plot, we can clearly see a strong deviation between the majority genotype cells and the microchimeric cells (Fig 4 d) indicating clear evidence of microchimerism consistent with samples with ground truth such as in silico mixtures. We then show kidney transplant biopsy data at days 5 and 28 after transplantation (Fig 4 e,f). We show a t-SNE of the kidney data showing the expected cell type specific separation between patient and donor cells as we expect primarily immune cell types to be a mix of patient and donor derived in a background of donor derived kidney cells. We used viewmastR^17^ to annotate cell types,training the model on a reference dataset^18^ with known cell type labels, and found that early after transplantation, primarily monocytes begin to infiltrate along with a minority of macrophages, plasma cells, and T cells. After 28 days we see larger populations of a variety of immune cell types from the patient infiltrating the transplanted kidney. We also tested Cellector’s robustness to the amount of sequencing data by downsampling synthetically mixed hipsci cell line data for 10 minority cells and 100 minority cells (Supp Fig 1). Cellector maintains near perfect PPV and >90% sensitivity until the median UMI drops below 2000 where both begin to degrade quickly.

**Fig. 4:**
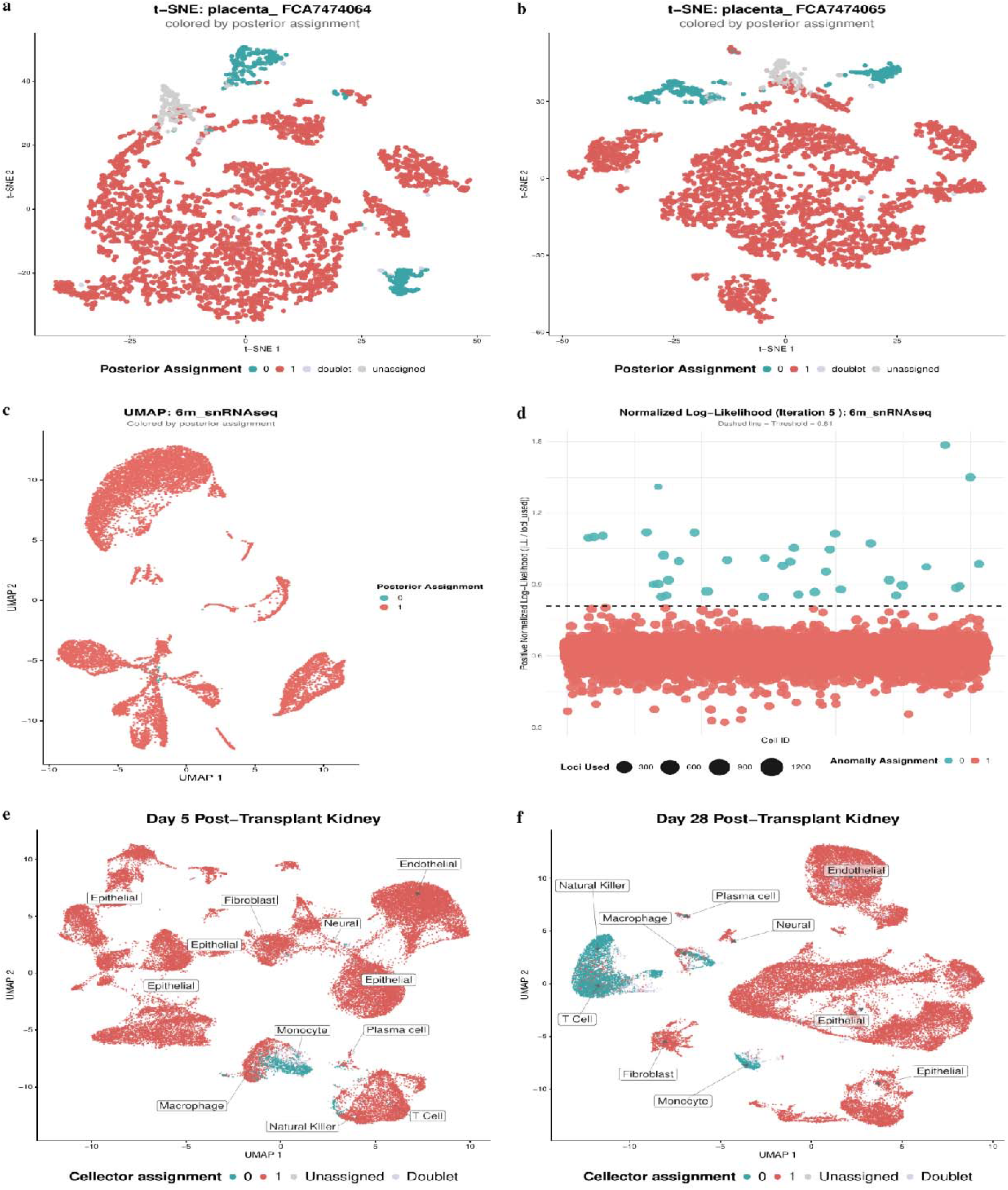
Demonstration on natural microchimeric maternal/fetal, neuronal, and kidney samples. **a,b** t-SNE plots of maternal/fetal cell samples colored by cellector assignment. **c** UMAP of neuronal sample with known microchimerism. **d** Anomaly likelihood plot of neuronal sample with known microchimerism. **e,f** UMAPs of kidney transplant samples at Day 5 and Day 28 colored by viewmastR and cellector assignments.

## Discussion

Here we have presented Cellector, a method for detecting exceedingly rare cells from foreign genotypes (including those from related individuals) in scRNAseq samples. This is vital for the study of microchimerism in general, but is specifically useful for the estimation of measurable residual disease in acute leukemias post hematopoietic cell transplant in which it could aid in the early detection of cancer relapse. If we can detect these events early, safer, less invasive and harmful therapeutics can be used. Cellector can also be used to better understand transplant biology and rejection from identifying invading immune cells into solid tumors and investigating their expression profiles. Additionally, cellector may be used to study natural microchimerism in maternal/fetal biology. This includes the evolution of the immune system in both the infant and mother and its effect on the viability of future pregnancies, the contribution to autoimmune diseases, and the development of the next generation’s immune system.

## Acknowledgements

None

## Funding

This work was supported by the NIH under the grant R01 CA289886-01; Subaward No. 0001188298.

## Conflict of interest

None

## Data availability

Cellector source code, singularity container, and documentation are released on GitHub: https://github.com/wheaton5/cellector.

The single-cell RNA-seq datasets used and analyzed in this study are publicly available. Figure 2 “unrelated” individuals was generated from an average across in-silico mixtures from hipsci samples available at the European Nucleotide Archive (ENA) under sample accessions ERS2630502–ERS2630507 corresponding to the cell lines euts, nufh, babz, oaqd, and ieki. Figure 2 “related” data will be made publicly available on GEO/SRA upon acceptance. Figure 4A and B maternal/fetal data are available at <u>E-MTAB-6701</u> with sample numbers FCA7474064 and FCA7474064. Figure 4C and D are derived from the neuronal sample GSM5138520 (6m_snRNAseq), available in GEO under accession number GSE168408. Single-nuclei RNA-seq and single-nuclei ATAC-seq data are deposited under the same accession^16^. Figure 4E and F are kidney transplant biopsy samples from accession GSE145927 in the Gene Expression Omnibus (GEO): GSM4339778 (Biopsy4_Day5; 5 days post-transplant, acute tubular injury without rejection) and GSM4339775 (Biopsy1_Day28; 28 days post-transplant, acute tubular injury without rejection)^19^

## Notes

https://github.com/wheaton5/cellector

### Competing Interest Statement

The authors have declared no competing interest.

