## supplement for "Cellector: A tool to detect foreign genotype cells in scRNAseq data with applications in leukemia and microchimerism"

**SUPPLEMENTARY**

1. **Downsampling experiments**

In order to assess how well Cellector performed with reduced data, we performed downsampling experiments to simulate lower amounts of sequencing and/or poor library complexity. For comparison, we included souporcell’s performance on the same data. We used the hipsci data set babz and added 10 and 100 cells from the hipsci sample euts_1 and then downsampled reads at downsample rates 2% to 98%. The starting median UMI per cell for these samples was 14,172 and 457 respectively. The number of total cells in the majority sample is 13708 representing a minority percentage of 0.073% and 0.073% for the 10 and 100 cell in silico spike-ins respectively. We see that with 10 minority cells, souporcell’s PPV is exceedingly low indicating a high false positive rate. But souporcell’s sensitivity hovers around 80% throughout the experiment indicating that the clustering did pick up on some genetic difference in addition to some random split in the data not associated with the sample differences but which optimized the loss function nonetheless (unwanted behavior). Cellector’s PPV remains at 100% until 98% dçownsampling representing a median UMI per cell of 457. Also in the 10 minority cell mixture, Cellector’s sensitivity remains >90% until around 80% downsampling. With 100 minority cells spiked in, Cellector’s PPV remained at 100% until 98% downsampling and its sensitivity was above 85% until ~90% downsampling. In these experiments, souporcell was able to find the appropriate clustering on genotype/sample lines but exchanged PPV for high sensitivity dropping below 80% PPV when the downsampling exceeded 50%. These experiments confirm that Cellector maintains exceedingly high PPV with reasonably high sensitivity until very extreme conditions are seen.

2. **Cancer simulations**

In order to assess whether Cellector could be used to detect cancer cells, we simulated a range of somatic mutations in a range of minority cells to see how well Cellector could identify them. We generated synthetic tumor datasets by introducing artificial somatic mutations into BAM files at predefined tumor cell fractions (2.5%, 5%, 10%, and 20%). We then chose different number of loci (6,250; 8,000; 12,500; 25,000; 50,000; 100,000; and 200,000) for which there was sequencing data, resulting in 35 total tumor–normal simulation conditions. Mutated reads were merged with the original alignments to reconstruct finalized tumor BAM files, which were then processed through a standard somatic variant calling workflow using Strelka2 to identify high-confidence PASS variants. At these detected variant loci, Vartrix was used to extract single-cell reference and alternate allele count matrices. We show that under very high mutational burden (>8000 mutations in the exome), Cellector can find the cancerous cells when they make up 5% or more of the sample. Of note is that the PPV remained at 100% throughout these experiments indicating an extremely low false positive rate. In combination with cell type and transcriptional profile, this is a promising future direction for Cellector.
